# Integrating genotype and weather variables for soybean yield prediction using deep learning

**DOI:** 10.1101/331561

**Authors:** Johnathon Shook, Linjiang Wu, Tryambak Gangopadhyay, Baskar Ganapathysubramanian, Soumik Sarkar, Asheesh K. Singh

**Affiliations:** Department of Agronomy, Iowa State University, Ames, IA, 50011, USA; Department of Mechanical Engineering, Iowa State University, Ames, IA, 50011, USA

**Author notes:** Corresponding Authors, A.K. Singh, Department of Agronomy; S. Sarkar, Department of Mechanical Engineering, Iowa State University, Ames, IA, 50011, USA. Funding information Funding for this project was provided by Iowa Soybean Association (AKS), Monsanto Chair in Soybean Breeding (AKS), Plant Sciences Institute (SS, BG and AKS), USDA (SS, BG, AKS), NSF NRT (graduate fellowship to JS).

**Keywords:** machine/deep learning, *yield prediction*, genotype-environment

## Abstract

Realized performance of complex traits is dependent on both genetic and environmental factors, which can be difficult to dissect due to the requirement for multiple replications of many genotypes in diverse environmental conditions. To mediate these problems, we present a machine learning framework in soybean (Glycine max (L.) Merr.) to analyze historical performance records from Uniform Soybean Tests (UST) in North America, with an aim to dissect and predict genotype response in multiple envrionments leveraging pedigree and genomic relatedness measures along with weekly weather parameters. The ML framework of Long Short Term Memory - Recurrent Neural Networks works by isolating key weather events and genetic interactions which affect yield, seed oil, seed protein and maturity enabling prediction of genotypic responses in unseen environments. This approach presents an exciting avenue for genotype x environment studies and enables prediction based systems. Our approaches can be applied in plant breeding programs with multi-environment and multi-genotype data, to identify superior genotypes through selection for commercial release as well as for determining ideal locations for efficient performance testing.

## 1 INTRODUCTION

Soybean (*Glycine max* (L.) Merrill) has a long history of cultivation in North America, with the first reported production in Georgia in 1766, predating the formation of the United States [1]. Over the years, production has expanded as far west as Kansas-Colorado border, and has been grown from southern Texas and into Canada where there is currently considerable expansion, such as the addition of Saskatchewan as a soybean-producing province [2, 3]. Climatic resiliency is an important objective for plant breeder and for farmers who are interested to grow high yielding and stable cultivars. The climatic variability is associated with changes in temperature and rainfall events: patterns and magnitude. In addition to spatial variability, temporal variability of weather variables [4] is equally important and generally less understood or not included in yield prediction equations. Prediction of the effects of changing environments on performance is critical for producers to allow them to make informed marketing decisions of their crops, optimize production by potentially shifting regional production to reflect the relative economic performance of different crops, and can also help breeders to compare results over multiple years to gain information about how experimental (pre-commercial) varieties will likely perform in a target environment [5].

North American annual soybean yield trials (known as Uniform Soybean Tests (UST)) have been coordinated through the United States Department of Agriculture (USDA) between public breeders in university and government settings in the United States and Canada since 1941 [6, 7]. These trials are used to evaluate current (as checks) and pre-commercial varieties in multiple environments within their range of adaptation. Participating programs measure a number of performance related traits, including seed yield, seed size, seed protein and oil content, plant height, lodging, days to maturity, and seed quality, with the completeness of records depending on labor availability and proximity of the testing site to a resource hub. Testing regions are split into the North and South, with the breakpoint occurring at Maturity Group (MG) IV. In a typical year, around 90 different environments are utilized for UST, ranging from southern Canada to Texas (MG 00-VIII), and are further split into cohorts based on maturity groups, stage (preliminary screenings; and Uniform for advanced stage testing), and, for the North region, conventional (non-genetically modified) vs Roundup Ready (RR). These tests are a valuable source of historical and current data for assimilation of genotype and environment variables for phenotype elucidation in a range of conditions.

In order to understand phenotype response and associated changes in performance for a given genotype between locations, it is necessary to first identify and explain differences between testing locations (i.e., environments). For example, performance for a given crop variety or hybrid will depend on management practices, such as row spacing, fertilization, planting date, and chemical control of disease or weeds, as well as on the genetics of the variety and the environment in which it is planted. This environmental component has been examined at small scales due to the labor required for managing large numbers of plots [8, 9]. Modern weather stations can provide sub-hourly measurements of multiple weather parameters known to be important, such as light intensity and quality, temperature, precipitation, and relative humidity. With the addition of each layer of additional characterization of the environment, less of the differences need to be ascribed to a generic “environmental” component, and can instead be examined individually and in combination with plant genetics.

Traditionally, crop growth models have been proposed to simulate and predict crop production in different scenarios including climate, genotype, soil and management factors [10]. These provide reasonable explanation on biophysical mechanisms and responses, however, these models have deficiencies related to input parameter estimation and prediction in complex and unforeseen circumstances [11]. Previous attempts at yield prediction across environments have relied on crop models generated by quantifying response in a limited number of lines while altering a single environmental variable, limiting the inference scope. Also, the ultimate deciphering of spatio-temporal-modularity separating long series crop season weather variables, and diverse locations or environments provide an additional challenge. Deep learning models can provide solutions to such complex data.

The traditional linear methods such as AutoRegressive Integrated Moving Average (ARIMA) have been used for time series forecasting problems, but these methods have many limitations which include specifying the number of past observations to be provided as input. Also, these methods mainly focus on univariate data and may not achieve high accuracy in multivariate time series forecasting problems. For time series prediction tasks, deep neural networks show robustness to noisy input and also have the capability to approximate arbitrary non-linear functions [12]. Despite their flexibility and power, deep neural networks can only be applied to problems whose inputs and targets can be sensibly encoded with vectors of fixed dimensionality and thus sequences pose a challenge for these networks [13]. Among deep learning for time series analysis, Long Short Term Memory Recurrent Neural Networks (LSTM-RNNs) are very useful as they can capture the long-term temporal dependencies in sequential data and have shown state-of-the-art results in various applications including off-line handwriting recognition [14], speech recognition, natural language processing, music generation, DNA sequence analysis, machine translation, and video activity recognition. In many such applications, the length of the input sequence may not be equal to the length of the output sequence. For example, in sentiment classification many-to-one LSTM architectures have to be implemented where feature vector representations of word embeddings are used at each time step as input.

With this setup, we propose an LSTM based framework for using multivariate time series to predict the yearly value of crop yield with 30 weeks (entire crop season) of data per year provided as input. The goal is to predict the trait response (seed yield, protein, oil, days to maturity, plant height, seed size) using 13 years of US and Canada UST data. Our proposed deep learning framework enables integration of large scale genotype and weather variable for such phenotype prediction for widespread applications in plant breeding (for making selections), field experimentation (for site selection), agriculture production (for yield estimation and crop selling decisions) along with many other applications.

## 2. MATERIALS AND METHODS

### 2.1 Preparation of Performance Records

Files from 2003-2015 were downloaded as PDFs[6, 7]. Pages not containing performance data were removed using Adobe Acrobat Pro DC. Using on-line utility Zamzar (Zamzar.com), all 26 PDFs from this period were converted to.xlsx files, with each tab corresponding to a single page in the file. In this way, the vast majority of tables were recovered with no errors or need for human translation. These tables were manually curated to align all performance records for a given genotype/location combination into a single row. Records which did not have yield data (due to a variety not being planted in a specific location, or dying prior to production of seed) were removed from the file. Following removal, data remained for a total of 104320 performance records over the 13 year period. In addition to yield per se, maturity date, height, lodging, seed size, seed quality, oil, and protein were compiled for analysis, as well as all available management information provided. 5609 unique genotypes are represented in the final dataset. After compilation, each performance record was imported to Python for future analysis.

### 2.2 Acquisition and sub-sampling of weather records

Daily weather records for all location/year combiniations were compiled based on the nearest weather station available from the 25km grid from Weather.com. These results were then down sampled to include maximum, minimum, and average conditions on a weekly time frame throughout the growing season (defined April 1st through November 31) and appended to the performance record data frame. The weather variables, maturity group, latitude/longitude, and year were then run through a LSTM model to identify which weather parameters were related to the observed differences in performance. Genotypes were then clustered based on the organization which bred them in order to provide some control over relatedness, to help improve the model performance. Models were tested for oil, protein, lodging, plant height, and yield. The original PDFs, including methods used for all measurements, can be found here [6] for the North, and here [7] for the South.

### 2.3 Genotype Clustering

Application of the model for specific genotypes rather than mean location yield across genotypes requires the inclusion of genotype-specific criteria. Due to the nature of the UST program, most of the genotypes tested in this period do not have marker data available, and many may have been completely discarded after testing, preventing the use of a G matrix. To circumvent these restrictions, we applied an initial clustering based on the organization which submitted each line to the UST. Commercial checks, which are included to provide a comparison to genotypes currently grown by farmers, were lumped into a single “Commercial” cluster, while genotypes developed by university or public research organizations were grouped by developer.

### 2.4 LSTM for Multivariate Time Series Prediction

Recurrent Neural Networks (RNN) are capable of explicitly capturing temporal/sequential correlations and dependencies in time series data. Efficient learning of the temporal dependencies leads to highly accurate predictions and forecasting, often outperforming static networks [15]. Typically, as most deep neural network training, deep RNNs are trained using the error backpropagation algorithm. However, the propagation of error gradients through the latent layers and unrolled temporal layers may have various issues such as the vanishing gradient problem. Therefore, gradient descent of an error criterion may be inadequate to train RNNs especially for tasks involving long-term dependencies [16]. Standard RNNs fail to learn in the presence of time lags greater than 5-10 discrete time steps between relevant input events and target signals [17].

Long short-term memory (LSTM) is a novel RNN architecture designed to overcome the error back flow problems [18]. By using input, output and forget gates to prevent the memory contents being perturbed by irrelevant inputs and outputs, LSTM networks have the ability in learning long range correlations in a sequence- the LSTM networks obviate the need for a pre-specified time window and are capable of accurately modeling complex multivariate sequences [19]. LSTM recurrent neural networks can therefore be effectively used for prediction tasks involving multivariate time series data as input.

LSTM networks are effective in capturing long term dependencies when the the gap between the relevant information and the point where it is needed becomes very large. The cell state in a LSTM block can allow the information to just flow along it unchanged and information can be added to or removed from the cell state, carefully regulated by structures called gates. The forget gate decides what information to be removed from the cell state. Forget gates naturally permit LSTM to learn local self-resets of memory contents that have become irrelevant [17]. The forget gate and output activation function are the most critical components of the LSTM block and removing any of them significantly impairs performance [20]. The input gate decides what new information is to be stored in the cell state. Also, the old cell state needs to be updated to a new cell state.

We use a many-to-one LSTM model where multivariate input in multiple time steps is used to predict a single value. The details of the input variables are shown in Figure 2. The problem formulation involves framing the dataset as a supervised learning problem and we are aiming to predict the yield for a particular year with the given input variables. All features are normalized before splitting the dataset into two sets- training set to train our proposed LSTM model and test set - to test the performance of the model. An error score can aid in evaluating the performance of the model. We calculate the Root Mean Square Error (RMSE) after inverting the applied scaling to have forecasts and the actual values in the original scale.

**FIGURE 1.**
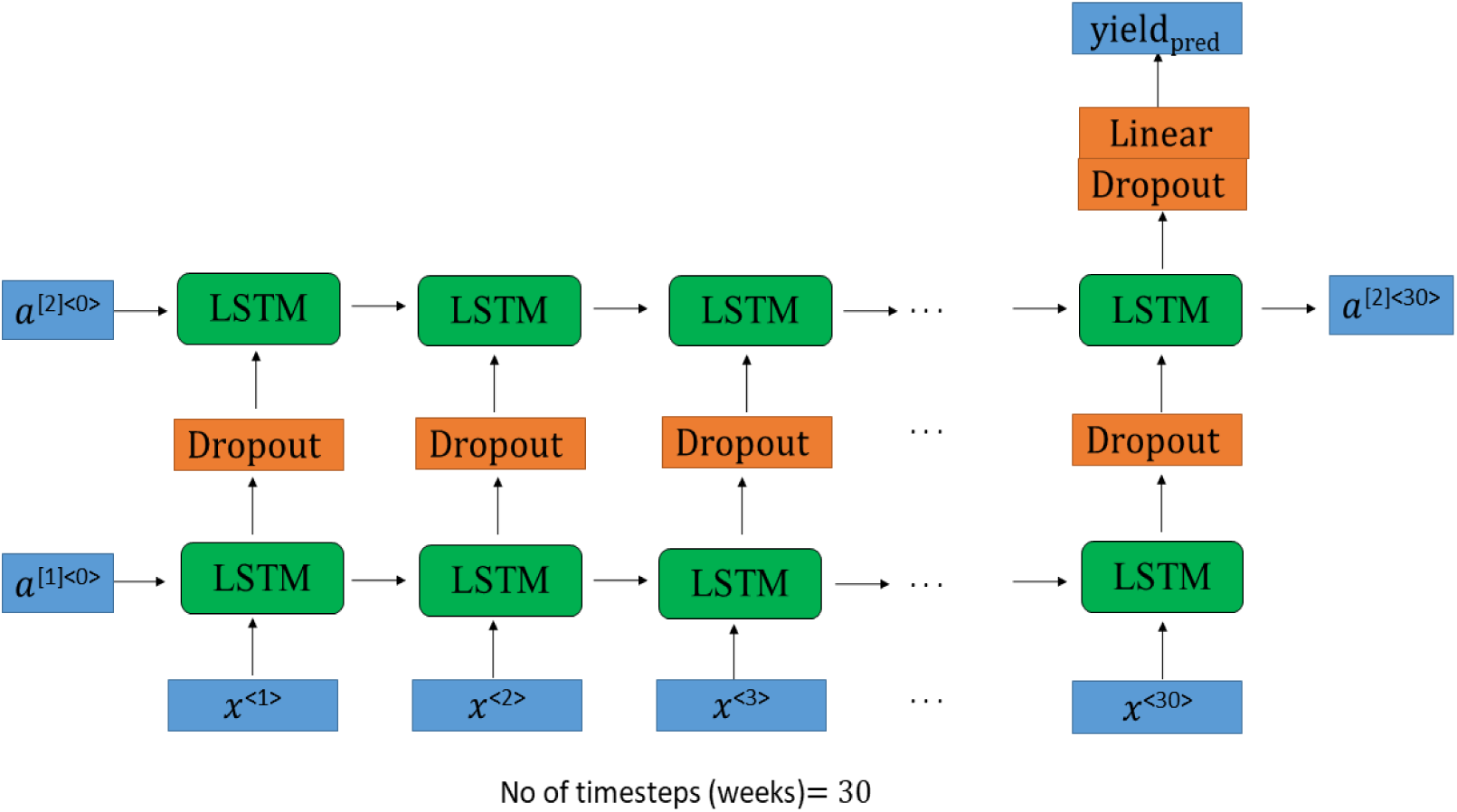
Many-to-one LSTM network architecture is implemented for 30 time steps (weeks). The output from the first LSTM layer is a batch of sequences propagated through another layer. The output from this LSTM layer is a single hidden state, not a batch of sequences. We use dropout regularization method after each LSTM layer to prevent overfitting.

**FIGURE 2.**
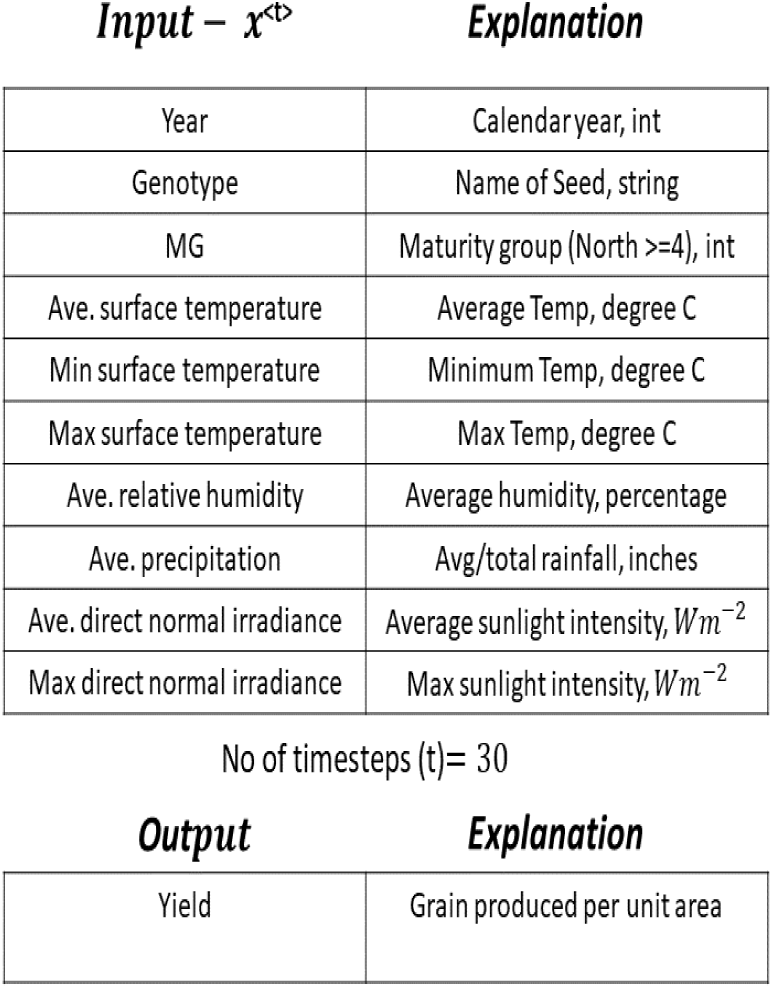
Multivariate input at each time step of the LSTM network and output variable is the yield

## 3 RESULTS

Initial results using only weather variables achieve a 68% accuracy in yield prediction in specific environments based on an 80/20 holdout dataset. The standard error of predictions was 10 bushels/acre. Improved performance was achieved by clustering genotypes by its originating institution and incorporating MG, GPS coordinates, and year. After these effects were included in the model, an accuracy of 82% was achieved. The RMSE of this model is 8.52. Figure 4 shows the yield prediction performance for 100 randomly selected test samples along with corresponding ground truth. It is evident from the plot that the model captures the general trend quite well while suffering in certain (rare and)extreme cases (yield too high or too low) which is a typical observation for most statistical models. The correlation plot considering all the test samples is demonstrated in Figure 3. Selections made based on model results, therefore, tend to correctly rank performance within a location, but predict the range of performance in a tighter band. Applying the prediction equation generated to each environment within the selected target population of environments (TPE) can be accurately used for prediction of the overall best yielding genotypes with reduced pre-commercial testing (locations and years)

**FIGURE 3.**
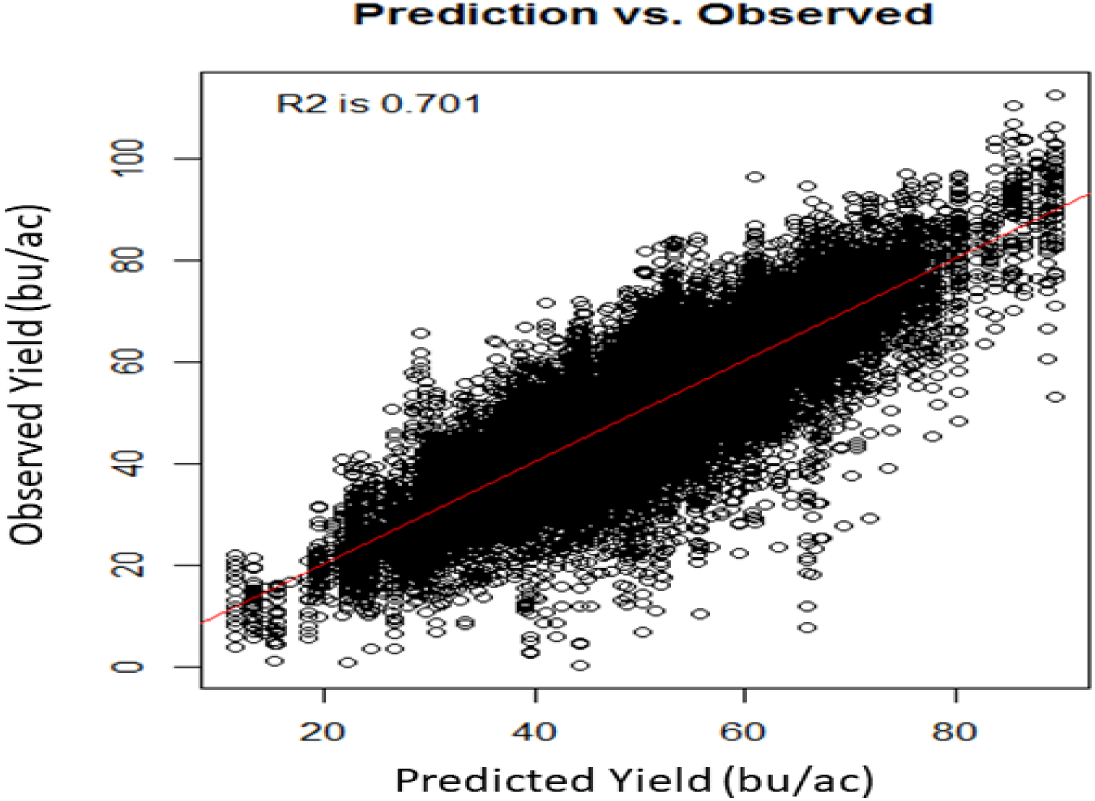
The correlation plot for all 20,000 test samples.

**FIGURE 4.**
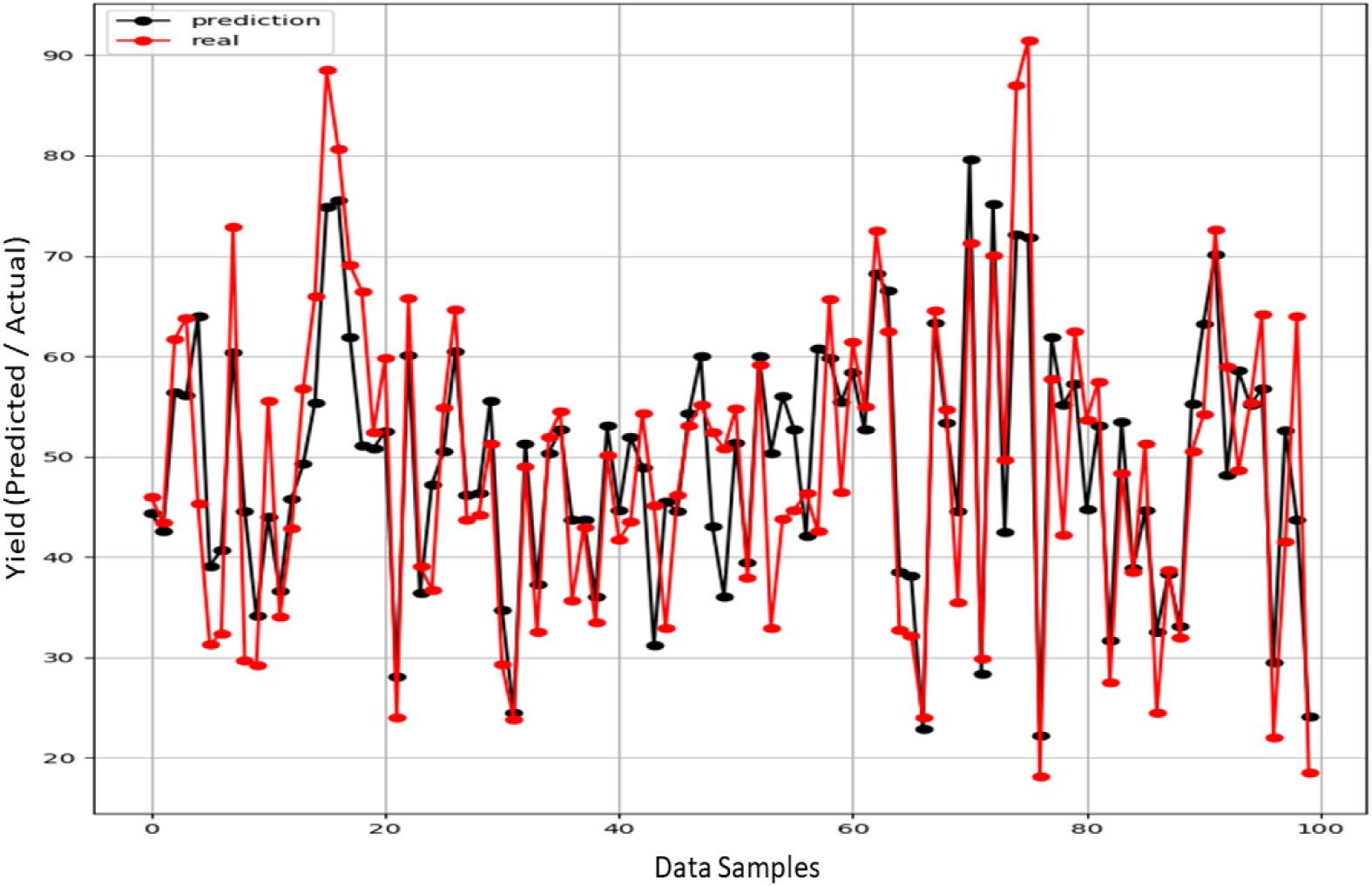
Comparison of Actual and Predicted Yield for 100 data samples selected randomly from the entire set of 20,000 test samples.

## 4 DISCUSSION

Deep LSTM is an efficient modeling scheme to analyze soybean crop growth interaction with the weather. For example, differences in the timing of extreme heat events, as well as drought periods, would affect soybean plants in various ways depending on the stage of development the plant is in. Heat stress during flowering, for example, is particularly damaging, while heat in vegetative stages of development may not produce significant detriment to harvested yield [21]. In such scenarios a live modelling approach presenting in this work using LSTM-RNN will be robust to incorporate these weather changes and adjust performance predictions accordingly.

Yield testing does not give a definite performance of an individual line in an environment, but rather an approximation [22]. Field variability, small plot size, low replication number, and error inherent in combine yield monitor are just a few examples of factors that increase the range of uncertainty, and can be reflected in the reported coefficient of variation (CV). By training a model with a smaller subset, one which utilizes only fields with small coefficients of variation (CVs), higher accuracy can be expected. This may be necessary for future attempts to dissect the genetic causes of differential response to weather; however, deep learning models can perform reasonably well if large data (even if heterogeous) is inputted. Use of a 13 year range may mean that some varieties that were tested early on may be direct parents of later tested varieties, further contributing to the data skewedness, and signifies the importance of integrating relationship matrix. Additional information that may improve the results of this approach are the inclusion of any supplemental irrigation provided, soil fertility levels, disease pressure and resistance levels, and direct genetic markers for the tested varieties, all of which would further strengthen predictive ability. With this data, it may become possible to identify QTL or even underlying genes which condition improved tolerance to different weather stresses, or that affect yield differences in normal environments, as well as to increase the model performance significantly [23]. These QTL can be immediately incorporated into breeding programs, with limited or no yield drag expected due to the improved nature of the lines being tested here.

Yield prediction based on weather records can have significant effects on the economies of agricultural states, with impacts on daily lives through food prices. We establish the potential for use of a long short-term memory method for yield prediction to allow models to account for temporal differences in the occurrence of weather events. Predictions using this system can be made reasonably accurate due to the large amount of training data made available through mining of historical records. Future implementations may be expanded to include pedigree or marker data, additional factors such as preceeding crop, row spacing, planting date, soil texture, or additional temporal data in the forms of soil sensor measurements or by using remote sensing.

## ACKNOWLEDGEMENTS

The authors would like to thank Vikas Chawla for his assistance with querying weather data for this project.

